# High-resolution population-specific recombination rates and their effect on phasing and genotype imputation

**DOI:** 10.1101/2020.05.20.106831

**Authors:** Shabbeer Hassan, Ida Surakka, Marja-Riitta Taskinen, Veikko Salomaa, Aarno Palotie, Maija Wessman, Taru Tukiainen, Matti Pirinen, Priit Palta, Samuli Ripatti

## Abstract

Founder population size, demographic changes (eg. population bottlenecks or rapid expansion) can lead to variation in recombination rates across different populations. Previous research has shown that using population-specific reference panels has a significant effect on downstream population genomic analysis like haplotype phasing, genotype imputation and association, especially in the context of population isolates. Here, we developed a high-resolution recombination rate mapping at 10kb and 50kb scale using high-coverage (20-30x) whole-genome sequenced 55 family trios from Finland and compared it to recombination rates of non-Finnish Europeans (NFE). We tested the downstream effects of the population-specific recombination rates in statistical phasing and genotype imputation in Finns as compared to the same analyses performed by using the NFE-based recombination rates. We found that Finnish recombination rates have a moderately high correlation (Spearman’s ρ =0.67-0.79) with non-Finnish Europeans, although on average (across all autosomal chromosomes), Finnish rates (2.268±0.4209 cM/Mb) are 12-14% lower than NFE (2.641±0.5032 cM/Mb). Finnish recombination map was found to have no significant effect in haplotype phasing accuracy (switch error rates ~ 2%) and average imputation concordance rates (97-98% for common, 92-96% for low frequency and 78-90% for rare variants). Our results suggest that downstream population genomic analyses like haplotype phasing and genotype imputation mostly depend on population-specific contexts like appropriate reference panels and their sample size, but not on population-specific recombination maps or effective population sizes. Currently, available HapMap recombination maps seem robust for population-specific phasing and imputation pipelines, even in the context of relatively isolated populations like Finland.

## 1. Introduction

Recombination is not uniform across the human genome with large areas having lower recombination rates, so-called ‘coldspots’, which are then interspersed by shorter regions marked by a high recombinational activity called ‘hotspots’ [1]. With long chunks of human genome existing in high linkage disequilibrium, LD [2], and organised in the form of ‘haplotype blocks’, the ‘coldspots’ tend to coincide with such regions of high LD [3].

Direct estimation methods of recombination are quite time-consuming, and evidence has suggested that they do not easily scale up to genome-wide, fine-scale recombinational variation estimation [4]. A less time-consuming but computationally intensive alternative is to use the LD patterns surrounding the SNPs [5]. Such methods have been used in the past decade or so, to create fine-scale recombination maps [6]. Besides the International HapMap project that focused on capturing common variants and haplotypes in diverse populations, international WGS-based collaborations like 1000 Genomes Project, provided genetic variation data for 20 worldwide populations [7]. This led to further refinement of the recombination maps coupled with methodological advances of using coalescent methods for recombination rate [8, 9].

With the rise of international collaborative projects, it was realised that founder populations can often have very unique LD patterns [10], subsequently also displaying unique increased genetics-driven health risks [11], suggesting that population-specific reference datasets should be used to leverage the LD patterns to better detect disease variants in downstream genetic analysis [12]. Genomic analysis methods like haplotype phasing and imputing genotypes require recombination maps and other population genetic parameters as input to obtain optimal results [13, 14, 15, 16]

In theis study, we set to test this by 1) estimating recombination rates along the genome in Finnish population using ~55 families of whole-genome sequenced (20-30x) Finns, 2) comparing these rates to some other European populations, and 3) comparing the effect of using Finnish recombination rate estimates and cosmopolitan estimates in phasing and imputation errors in Finnish samples.

## 2. Materials & Methods

### 2.1 Datasets used

#### Finnish Migraine Families Collection

Whole-genome sequenced trios (n = 55) consisting of the parent-offspring combination were drawn from a large Finnish migraine families collection consisting of 1,589 families totalling 8,319 individuals [17]. The trios were used for the recombination map construction using LDHAT version 2. The families were collected over 25 years from various headache clinics in Finland (Helsinki, Turku, Jyväskylä, Tampere, Kemi, and Kuopio) and via advertisements in the national migraine patient organisation web page (https://migreeni.org/). The families consist of different pedigree sizes from small to large (1-5+ individuals). Of the 8319 individuals, 5317 have a confirmed migraine diagnosis based on the third edition of the established International Classification for Headache Disorders (ICHD-3) criteria [18].

#### EUFAM cohort

To check the phasing accuracy of our Finnish recombination map, we used an independently sourced 49 trios from the European Multicenter Study on Familial Dyslipidemias in Patients with Premature Coronary Heart Disease (EUFAM). Finnish familial combined hyperlipidemia (FCH) families were identified from patients initially admitted to hospitals with premature cardiovascular heart disease (CHD) diagnosis who also had elevated levels of total cholesterol (TC), triglycerides (TG) or both in the ≥ 90th Finnish population percentile. Those families who had at least one additional first-degree relative also affected with hyperlipidemia were also included in the study apart from individuals with elevated levels of TG. [19, 20, 21].

#### FINRISK cohort

The imputation accuracy of the Finnish and previously published HapMap based recombination maps [8, 9] was subsequently tested on an independent FINRISK CoreExome chip dataset consisting of 10,481 individuals derived from the national-level FINRISK cohort. Primarily, it comprises of respondents of representative, cross-sectional population surveys that are conducted once every 5 years since 1972 to get a national assessment of various risk factors of chronic diseases and other health behaviours among the working-age population drawn from 3 to 4 major cities in Finland [22].

#### FINNISH reference panel cohort

The whole-genome sequenced samples used were obtained from PCR-free methods and PCR-amplified methods, which was followed by sequencing on a Illumina HiSeq X platform with a mean depth of ~30×. The obtained reads were then aligned to the GRCh37 (hg19) human reference genome assembly using BWA-MEM. Best practice guidelines from Genome Analysis Toolkit (GATK) were used to process the BAM files and variant calling. Several criteria were used in this stage for sample exclusion: relatedness (identity-by-descent (IBD) > 0.1), sex mismatches, among several others. Furthermore, samples were filtered based on other criteria such as: non-reference variants, singletons, heterozygous/homozygous variants ratio, insertion/deletion ratio for novel indels, insertion/deletion ratio for indels observed in dbSNP, and transition/transversion ratio.

After this stage, some exclusion criteria were applied to set some variants as missing: GQ < 20, phred-scaled genotype likelihood of reference allele < 20 for heterozygous and homozygous variant calls, and allele balance <0.2 or >0.8 for heterozygous calls. A truth sensitivity percentage threshold of 99.8% for SNVs and of 99.9% for indels was used based on the GATK Variant Quality Score Recalibration (VQSR) to filter variants with, quality by depth (QoD) < 2 for SNVs and < 3 for indels, call rate < 90%, and Hardy-Weinberg equilibrium (HWE) p-value < 1×10-9. Some other variants like monomorphic, multi-allelic and low-complexity regions [23] were further excluded. The final reference dataset used in this study for imputation consisted of high coverage (20-30x) whole-genome sequence-based reference panel of 2690 individuals from the SISu project (Sequencing Initiative Suomi, http://www.sisuproject.fi/, [24]).

### 2.2 Recombination map construction

Coalescent-based fine-scale recombination map construction [8] is greatly eased by using trios which provide more accurate haplotype phasing resolution [25]. Hence, we used trio data (n=55, 110 independent parents) from the Finnish Migraine Families Cohort described above. These were filtered primarily using VCFtools [26] and custom R scripts. Firstly, sites were thinned with within 15bp of each other such that only one site remained followed by a filtering step of removing variants with a minor allele frequency of <5% [27]. The resultant data were then phased using family-aware method of SHAPEIT [28] using the standard HapMap recombination map [8, 9], which was then split into segments of ~10000 SNPs with a 1000 SNP overhang on each side of the segments. LDhat version 2 was run for 10^7^ iterations with a block penalty of 5, every 5000 iterations of them of which the first 10% observations were discarded [8, 29]. The CEU based maps, used here for comparison, were obtained similarly using LDhat [29].

However, LDHat is computationally intensive, and calculations suggest that the 1000 Genomes OMNI data set [30] would be too much computationally intensive to complete [31], hence limiting the maximum number of haplotypes which could be used.

To overcome this and make the recombination map independent of the underlying methodology, we used a machine learning method implemented in FastEPRR [31, 32]. It supports the use of larger sample sizes, than LDHat and the recombination estimates for sample sizes > 50, yields smaller variance than LDHat based estimates [31]. The method was then applied to each autosome with overlapping sliding windows (*i.e.*, window size, 50 kb and step length, 25 kb) under default settings for diploid organisms. As seen in [31] both methods produce similar estimates, with only variance of the estimate of mean being different.

The output of LDHat and FastEPRR is in terms of population recombination rate (p) and to convert them into per-generational rate (r) used in phasing/imputation algorithms we used optimal effective population size values derived from our testing (as explained in the Supplementary Text). The estimates from LDHat and FastEPRR were then averaged, to obtain a new combined estimate with the lowest variance amongst all the three [31].

### 2.3 Phasing and imputation accuracy

To test whether the usage of different recombination maps affects the efficiency of haplotype phasing and imputation, we used the aforesaid Finnish genotype data to evaluate: (i) switch error rates across all chromosomes and (ii) imputation concordance rates for chromosome 20.

#### 2.3.1 Phasing Accuracy

The gold standard method to estimate haplotype phasing accuracy is to count the number of switches (or recombination events) needed between the computationally phased dataset and the true haplotypes [33].The number of such switches divided by the number of all possible switches is called switch error rate (SER).

For testing the influence of recombination maps on phasing accuracy, we used three different recombination maps: HapMap, fine-scale Finnish recombination map and a constant background recombination rate (1cM/Mb), to phase the 55 offspring haplotypes without using any reference dataset. To check whether reference panels used during haplotype phasing made any impact on the switch error rates, we used the Finnish SISU based reference (n=2690), to check whether the size of the reference panel made any impact on the results in phasing the offspring’s haplotypes (Figure 1). The SER in the offspring’s phased haplotypes were then calculated by determining the true offspring haplotypes using data from the parents (98 individuals) with a custom script [34].

**Figure 1:**
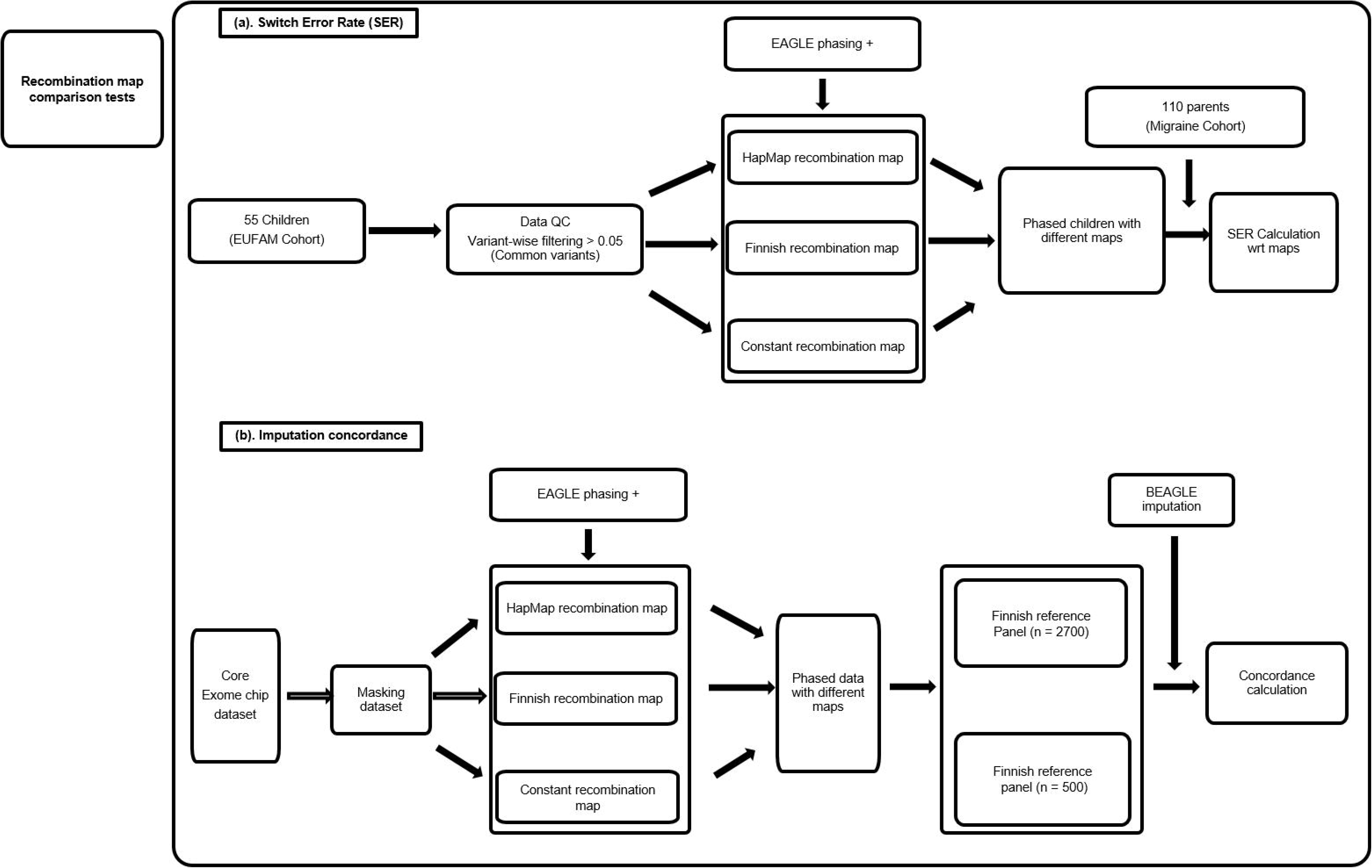
Flowchart overview of the analyses and comparisons performed.

#### 2.3.2 Imputation Accuracy

Imputation concordance was used as the metric for calculating the imputation accuracy. For this, we randomly masked FINRISK CoreExome chip data consisting of 10,480 individuals [22] from chromosome 20. To test the role of reference panel size in influencing the imputation accuracy in conjunction with varying the population genetics parameters, we imputed the masked dataset with BEAGLE (Browning *et al.*, 2016) using the Finnish reference panel (n = 2690). The concordance was then calculated between the imputed genotypes and the original masked variants. Masking was done by randomly removing ~10% of variants from the chip dataset.

The influence of recombination maps on imputation accuracy was checked by calculating the concordance values between imputed and original variants, using the Finnish reference panel in various combinations of recombination maps (constant rate, HapMap, Finnish map) during the imputation (Figure 1).

## 3. Results

### 3.1 Finnish recombination map and its comparison to the HapMap recombination map

The primary aim of our study was to derive a high-resolution genetic recombination map for Finland and use it for comparative tests in commonly used analyses like haplotype phasing and imputation. To derive a population-specific Finnish recombination map, we used the high-coverage WGS data and an average of different estimation methods (LDHat and FastEPRR). We used the Ne value of 10,000 derived from our extensive testing of different Ne values (See supplementary text) to get the per-generation recombination rates. The average recombination rates of Finnish population isolate depicted 12-14% lower values (autosome-wide average 2.268±0.4209 cM/Mb) for all chromosomes compared to CEU based maps (2.641±0.5032 cM/Mb) (Figure 2).

**Figure 2:**
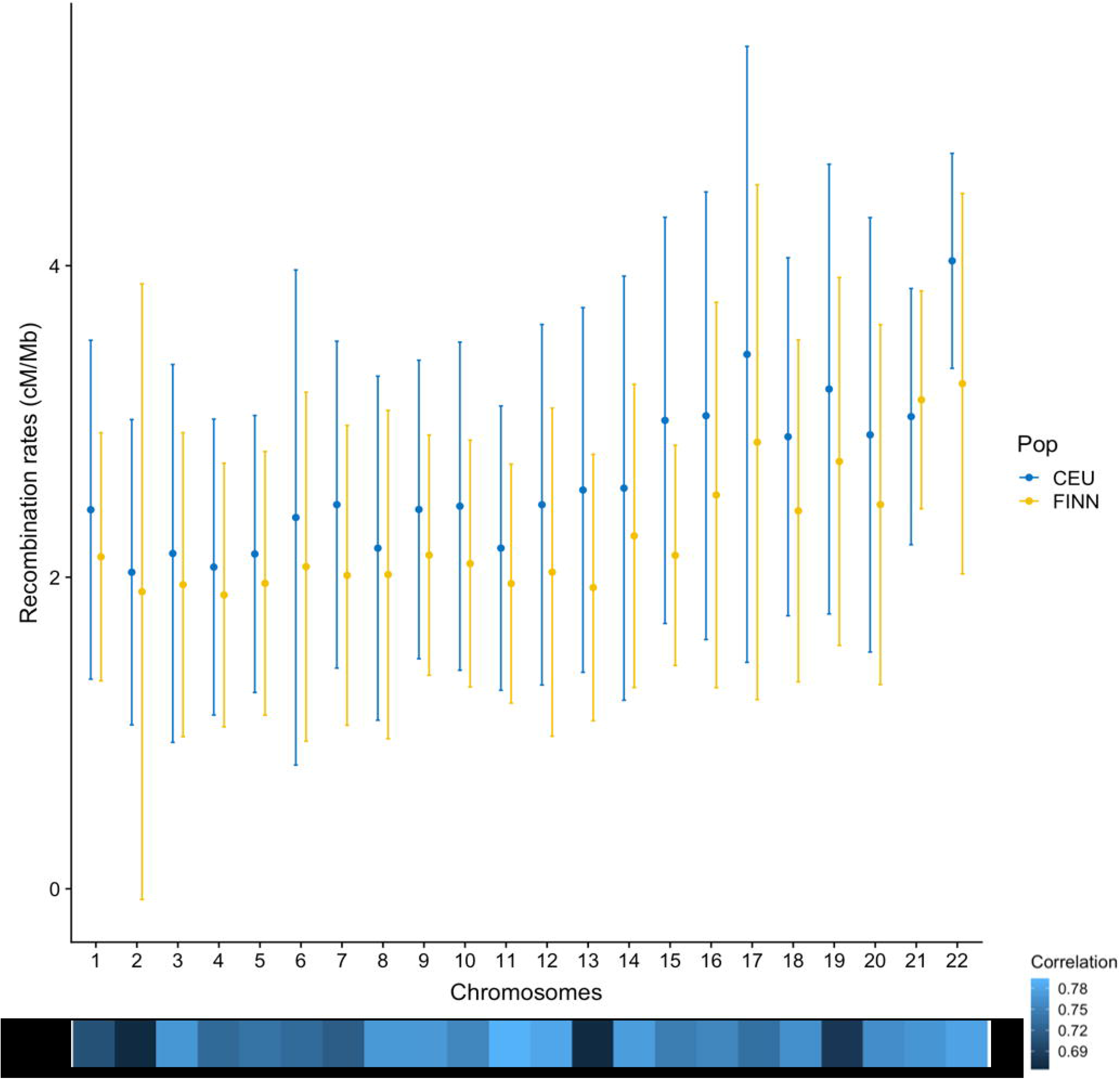
Average (± standard deviation) recombination rates of Finnish v/s CEU per autosome measured in cM/Mb and Correlation between Finnish and CEU recombination rates across all chromosomes. The comparisons are made for similar physical positions.

These differences in average recombination rates are reflected in the correlation values across all chromosomes (Spearman’s ρ ~ 0.67-0.79) between the developed Finnish map and HapMap based one (Figure 2). We also present a direct comparison between the two maps, of the recombination rates at 5Mb scales, which presents a similar visual pattern of rates across the genome (Supplementary Figure 1).

### 3.2 Effects of the population-specific recombinations map on haplotype phasing

Variation in population-specific recombination maps (and effective population sizes) can affect the downstream genomic analyses like haplotype phasing and imputation. We tested the Finnish map, HapMap map and a constant recombination rate map (1cM/Mb) to understand the effects of population-specific maps on downstream genomic analyses. The phasing accuracy was tested under two different conditions: using no additional reference panel and using an population-specific. SISu v2 reference panel (n= 2690) in phasing. We observed that, on average, SER ranged between 1.8-3.7% (Supplementary Figure 2) across the different chromosomes and recombination maps. We found statistically significant differences within both no-reference panel and the Finnish reference panel results (Kruskal Wallis, p-value = 5.3e-10 and 4.7e-10, respectively; Figure 3). The constant recombination map (1cM/Mb) had significantly higher SER values when compared to the Finnish map or the HapMap map (Figure 3) both when no reference panels were used (p-value = 2.9e-11 and 2.6e-09, respectively) and when the Finnish reference panel was used (p-value = 2.9e-11 and 9.5e-13, respectively). The choice of recombination maps mattered more when no reference panel was used (p-value = 0.0046), however when using the Finnish reference panel, the difference in SER was statistically insignificant (p-value = 0.25).

**Figure 3:**
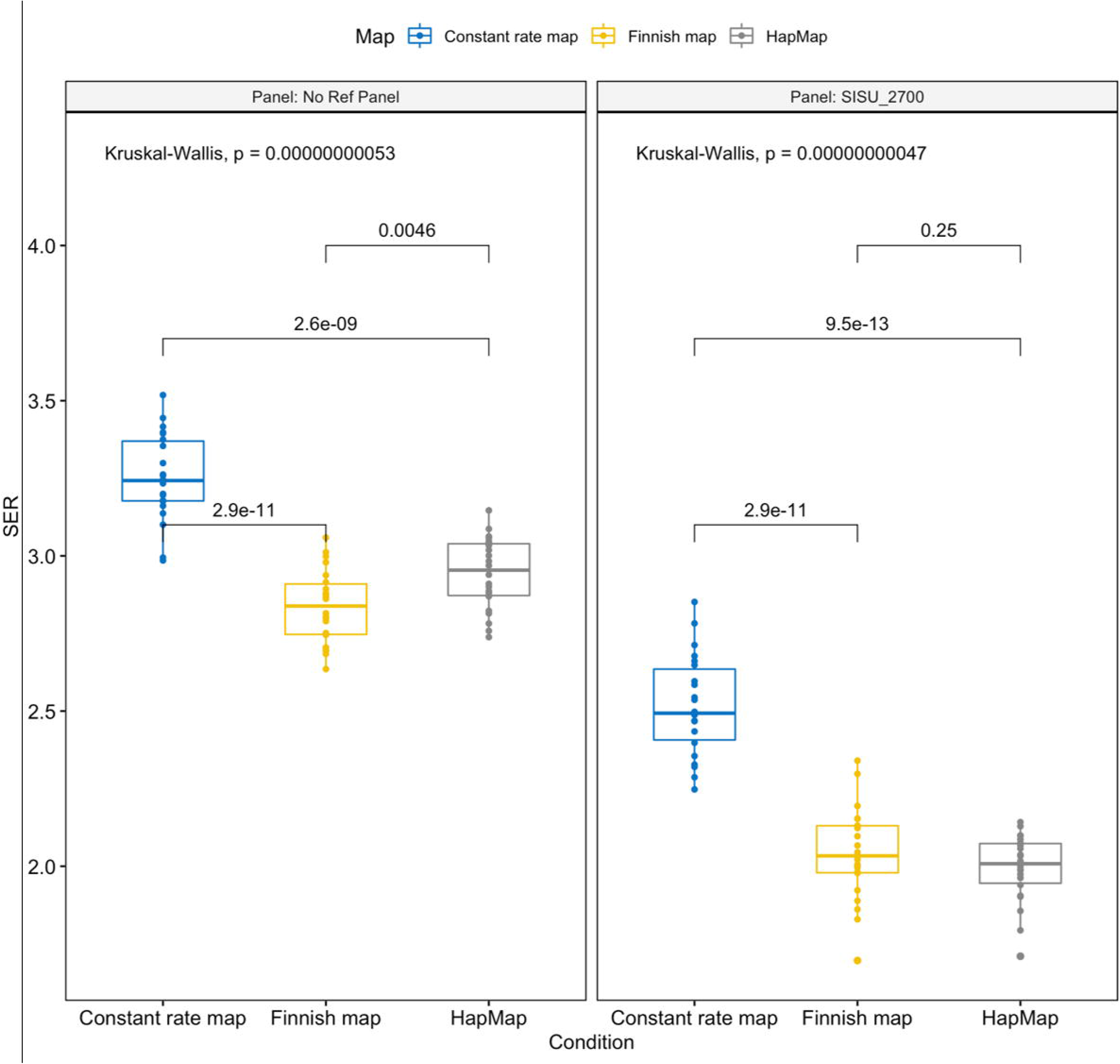
Statistical comparison of Switch Error Rates across all autosomes calculated for all children in the trios using different recombination maps with respect to different reference panel conditions (absent or present). The p-values are shown at the top of each panel from Kruskal Wallis ANOVA testing between panel groups and ones between boxplots for within-group comparisons.

### 3.3 Effects of the population-specific recombinations map on genotype imputation

Imputation accuracy was similarly tested using the reference panel under three different recombination map settings. We observed that when the imputation target dataset was phased and imputed using the Finnish reference panel (n=2690) irrespective of the population-specific recombination maps, it had a high imputation accuracy (overall concordance rate ~98%, Figure 4) across MAF bins (>0.1%). Though some differences in concordance rates are seen in for rare variants (MAF <0.1%). The concordance rate was lower when the test dataset was phased without reference panels (concordance rate 72~77%, Figure 5).

**Figure 4:**
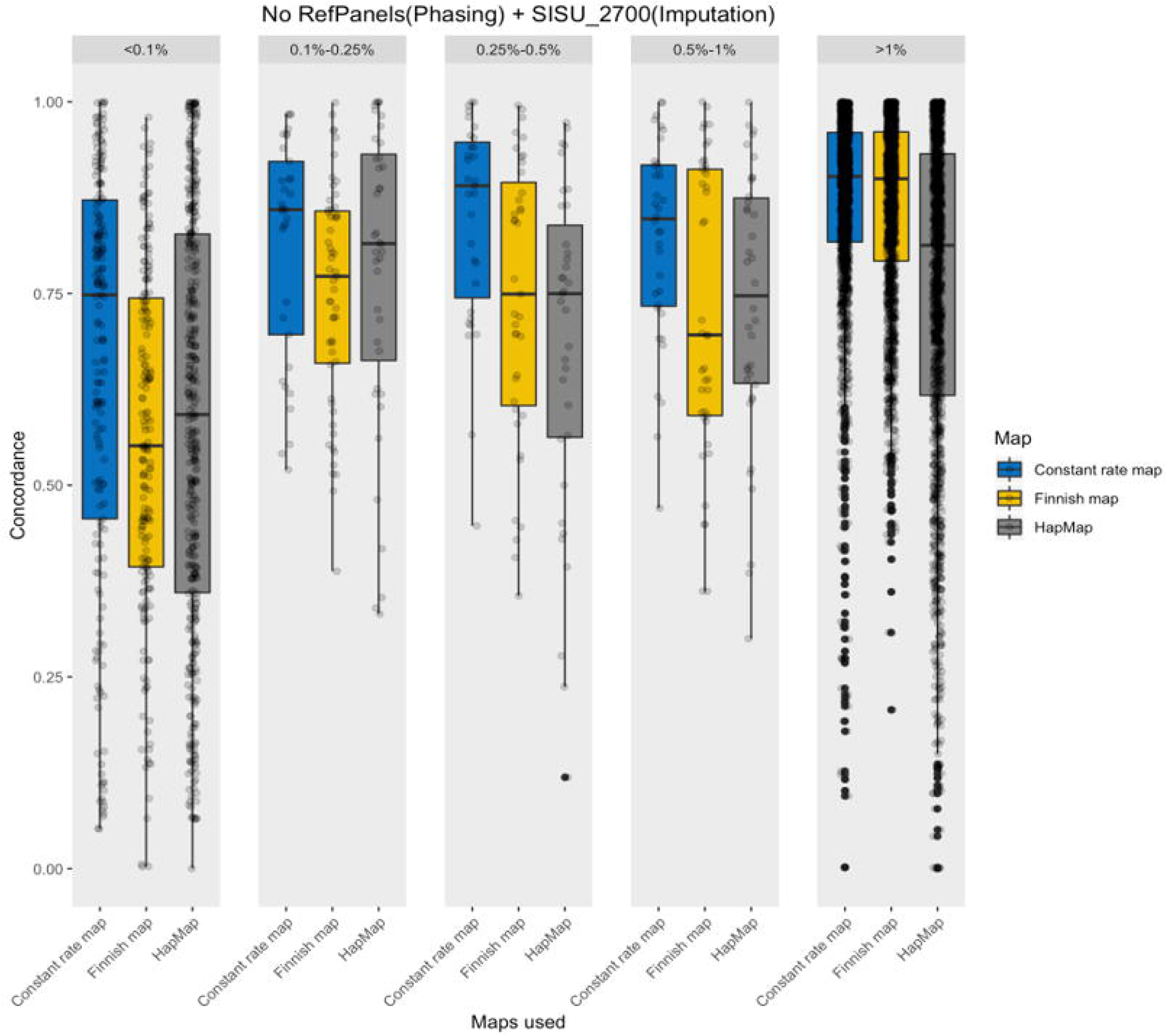
Comparison of Imputation Concordance across different Minor Allele Frequency (MAF) groups for a range of different recombination map combinations phased with NO reference panels

**Figure 5:**
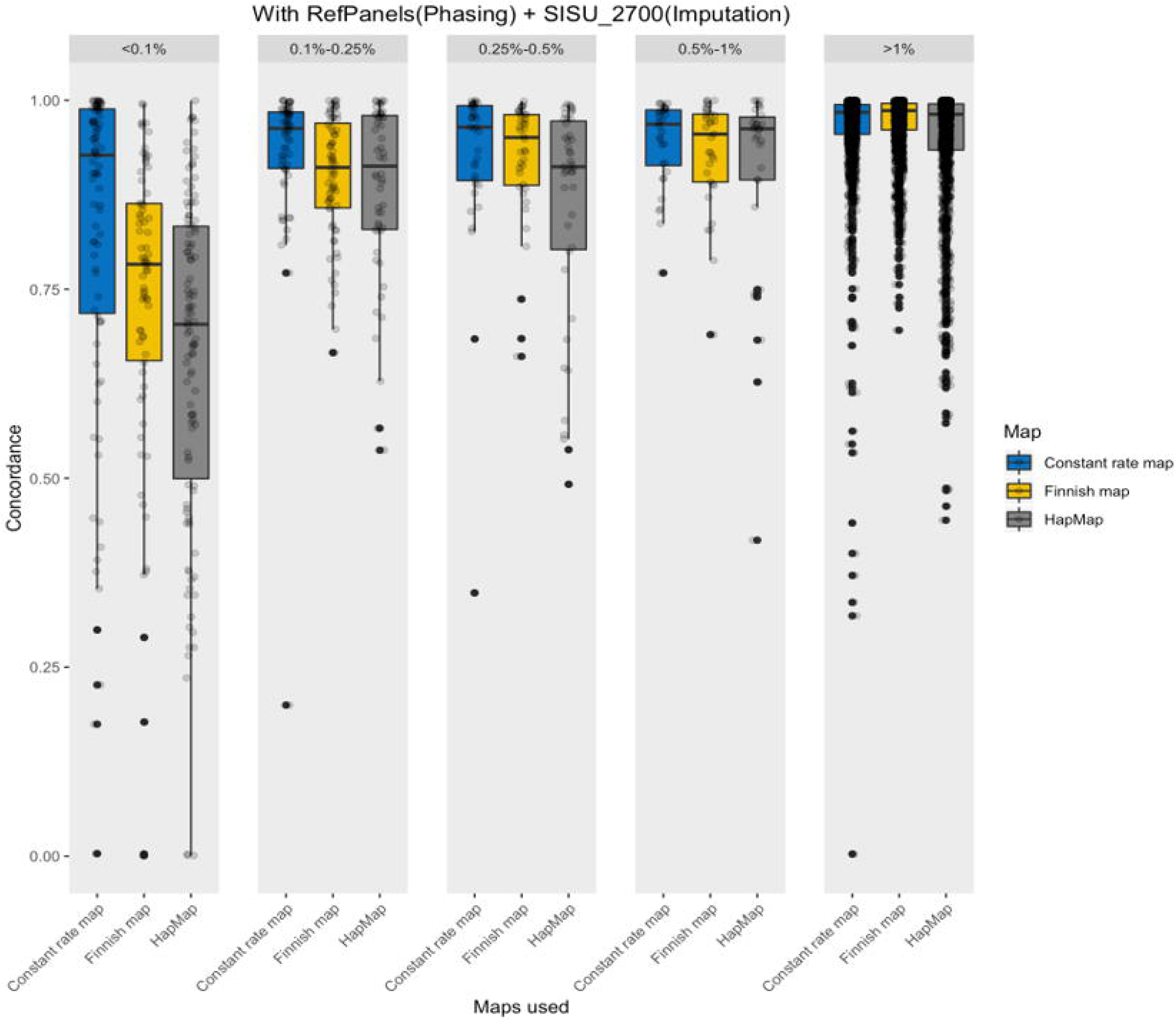
Comparison of Imputation Concordance across different Minor Allele Frequency (MAF) groups for a range of different recombination map combinations phased with reference panels.

## 4. Discussion

Population isolates like Finland, have had a divergent demographic history as compared to the outbred European populations, with a less historic migration, more fluctuating population sizes and higher incidences of bottleneck events and founder effects [35, 36] This unique demographic history then affects different population genetic parameters, like recombination rates [37]. It has been shown previously that using population-specific genomic reference panels augmented the accuracy of imputation accuracy leading to better mapping of diseases specific variants in GWAS [12]. Since recombination rates (in the form of recombination maps), features in much of the downstream genomic analyses’ methods like imputation and haplotype phasing [15, 34], we wanted to study their effect on downstream analyses.

Firstly, we characterised the Finnish recombination map using high-coverage (~30x) whole-genome sequencing (WGS) samples from large SISu v2 reference panel (n=2690). Previously used recombination maps hail from the HapMap and 1000Genomes projects which used sparse genotypic datasets or low-depth sequencing samples. This is a first attempt in creating a recombination map for Finland using population-specificWGS samples. We used two different methods in estimating the recombination rates, to achieve accurate estimates with lower variance [29,31]. In addition, we estimated effective population sizes using identity-by-descent (IBD) based methods [15] for both Finnish and CEU based datasets. The obtained recombination map was then used to test their role and importance in two selected downstream genomic analyses – haplotype phasing and imputation concordance. Since the recombination rate determination requires effective population size estimates, we also tested the role of varying effective population size on these two analyses (See Supplementary Text). The extensive testing of Ne yielded the estimate of 10,000 originally derived theoretically [38] and most used commonly for humans fits quite rightly for the recombination map.

The Finnish recombinational landscape when compared to the HapMap based map, showed, on average, a high degree of correlation across scales (10, 50kb and 5Mb), however, on average, Finnish recombination rates across chromosomes were found to be lower. Such moderate to high correlations (Figure 2) and similar recombinational landscape (Supplementary Figure 1) could be due to high sharing of recombinations in individuals from closely-related populations. The degree of dissimilarity in the population-level differences between Finnish and mainland Europeans in terms of recombination rates could be due to population-specific demographic processes like founder effects, bottleneck events and migration [39], or chromatin structure PRDM9 binding locations, for example [40]. And the broad similarity in terms of correlational structure seen here, reflects a shared ancestral origin of Finns and other mainland Europeans [41]. Other studies on population isolates like Iceland [9] have previously found a high degree of correlation with CEU based maps, albeit with substantial differences as seen here. Previous studies [42] have additionally explored the relationship between recombination rate differences between populations and allele frequency differences, with evidence suggesting that the differences between rates show the selection impact in the past 100,000 years since the out-of-Africa movement of humans.

As seen in previous studies, much of the downstream genomic analyses like getting more refined GWAS hits or, accurate copy number variants (CNV) imputation, can be highly improved with the addition/use of population-specific datasets [12]. To test this in the context of population-specific recombination maps, we used them to test the haplotype phasing and imputation accuracy and observed that despite large differences in the effective population sizes between populations, it did not affect the tested metrics. One possible explanation for the insignificant effect seen here is that the role of parameters like effective population size and recombination maps is to scale over the haplotypes for efficient coverage of the whole genome. However, when sufficiently large, population-specific genomic reference panels are available with tens of thousands of haplotypic combinations, such scaling over for specific populations, does not yield in substantial improvements. As we showed here, reference panel size could play an important role in the downstream genomic analyses and in most cases, the current practice of using the standard HapMap recombination map can be reasonably used. Another point of interest here is that the use of different Ne parameters during phasing/imputation might be redundant as we observed no change in the accuracy of our estimates on varying the Ne parameters. Similarly, when using population-specific recombination maps, we did not find any tangible benefits in using them over the current standard maps based on the HapMap data.

Our study suggests a couple of important points for future studies: (a) varying effective population size for downstream genomic analyses, such as phasing and imputation, might have a relatively small impact, and it might be better to use the default option of the particular software; (b) when available, it is beneficial to use a population-specific genomic reference panel as they increase the accuracy; (c) HapMap can be used for current downstream genomic analyses like haplotype phasing or genotype imputation in European-based populations. And, if need be, can be substituted for using population-specific maps, as the accuracy rates are quite similar to the population-based maps.

Though the sample used here is from a disease cohort but is nevertheless representative of Finland’s population and hence provides a reasonable recombination rate estimates. On the other hand, our reliance on disease cohorts could lead to minor variation in the resultant recombination. Though as we share a similar out-of-Africa origin, much of our history is shared and though biological differences in the recombinational landscape do exist between different populations, much of the downstream genomic analyses (haplotyping, imputation or, GWAS), might not be affected by recombination map or values of effective population size.

## Acknowledgements

We would like to thank Sari Kivikko and Huei-Yi Shen for management assistance. The FINRISK analyses were conducted using the THL biobank permission for project BB2015_55.1. We thank all study participants for their generous participation in the FINRISK study.

