## Supplementary Information for "High-resolution population-specific recombination rates and their effect on phasing and genotype imputation" (from another version of this preprint: version 2, 2020-05-23)

**Recombinational Landscape**

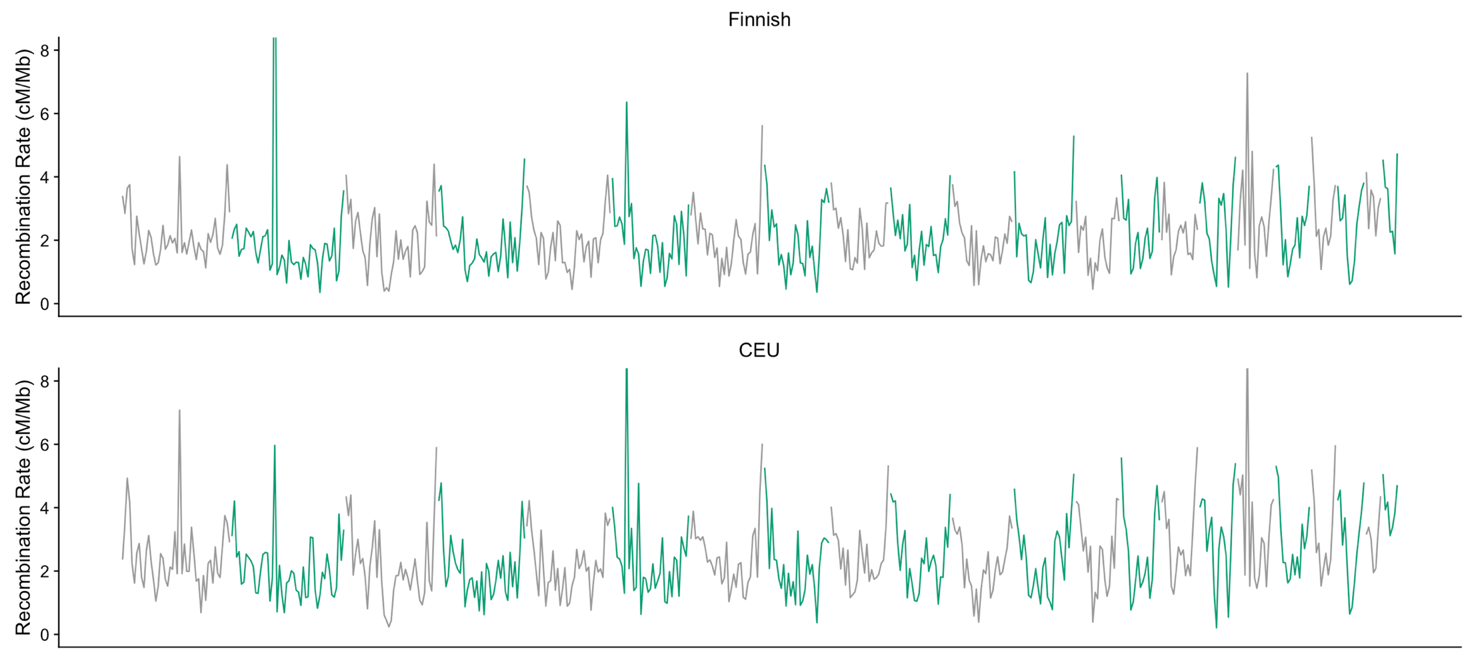

**Figure S1:** Comparison of Finnish and CEU recombination rates in 5Mb scale across all autosomes. The chromosomes are alternately coloured.

**Effective Population Size calculation**

To infer the historical effective population size, we used a non-parametric method IBDNe v. r1206 (Browning and Browning, 2015), based on a Wright–Fisher discrete-generation model. The input to IBDNe is a list of ancestry-specific identity-by-descent (IBD) segments obtained from IBDseq v.r1206 (Browning and Browning, 2015). We used the default settings to filter IBD segments with any breaks and short gaps in those segments were then removed with the merge-ibd-segments utility program (Browning and Browning, 2015). Using these segments, IBDNe was run with the default parameters to provide information on Ne for around 50 generations., wherein each generation was taken to be around 25 years.

Different populations with their unique demographic histories affect population genetic parameters used in genomic analyses. A prime example of this is the effective population size (Ne), and here we estimated Ne for 50 generations (~1250 years), from the present, utilising the IBD segments with IBDNe (Figure 4). We found that the effective population sizes (Ne) varied widely between Finland and rest of Europe (CEU) over the past 50 generations. The median Ne for Finland was found to be lower than CEU (4120 v/s 525,000), 50 generations (~1250 years) ago and current values (10^6^ v/s 10^8^) are still quite different. Taking the census data from 1750 onwards, we see that Finnish population increased from 420,000 in 1750 to 4.4 million in 1960, witnessing a 10-fold increase (Statistics Finland, http://www.stat.fi/til/vaerak/tau_en.html). Ne which is usually 1/3^rd^ of the census size follows the same pattern (44800 in 1750 to 1.2 million in 1960).

The change in recent Ne has been exponential with noticeable past bottleneck events in both populations highlighting the role of past demographic processes (migration, expansion, founder effects or bottlenecks) (Wang *et al.,* 2016)

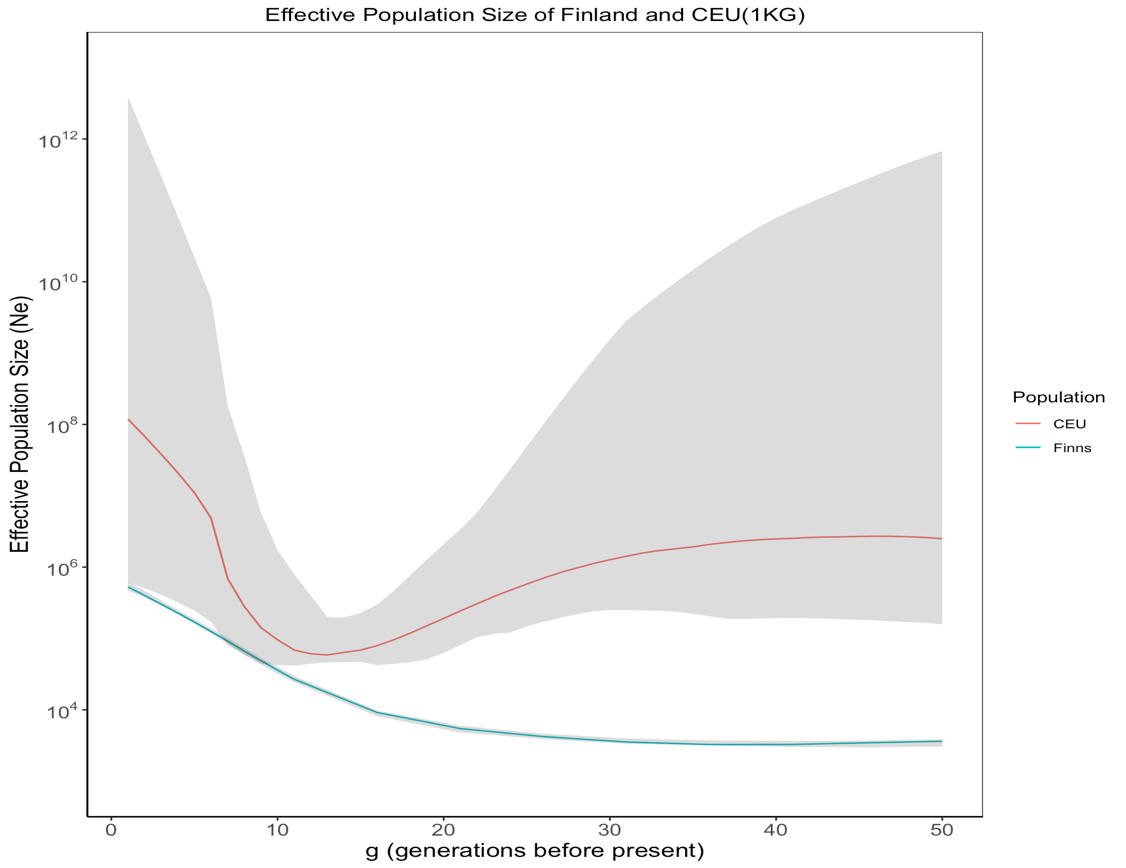

**Figure S2**: Comparison of effective population sizes (with shaded confidence intervals) over time up to 50 generations between Finland (SISU Reference Panel, n=2690) and CEU (Utah residents (CEPH) with Northern and Western European ancestry) dataset from 1000Genomes Release 3. Each generational period is considered to be 25 years.

**Effective population size comparisons**

For testing the influence of effective population size, we used the HapMap recombination map (Myers *et al.,* 2005) and phased the offspring (49 individuals) from the 49 trios (147 individuals) of EUFAM cohort by a popular phasing software, EAGLE (Loh *et al.,* 2016). The following values of Ne were used for each phasing run: 5000, 10000, 15000, 20000, 25000, 30000, 100000. For testing the influence of effective population size, we used the Finnish reference dataset (n = 2690) to impute the masked dataset while varying the effective population size (Ne) parameter from 1000 to 1 million and the concordance calculated as mentioned before. The flowchart for the comparisons is depicted below (Figure S3).

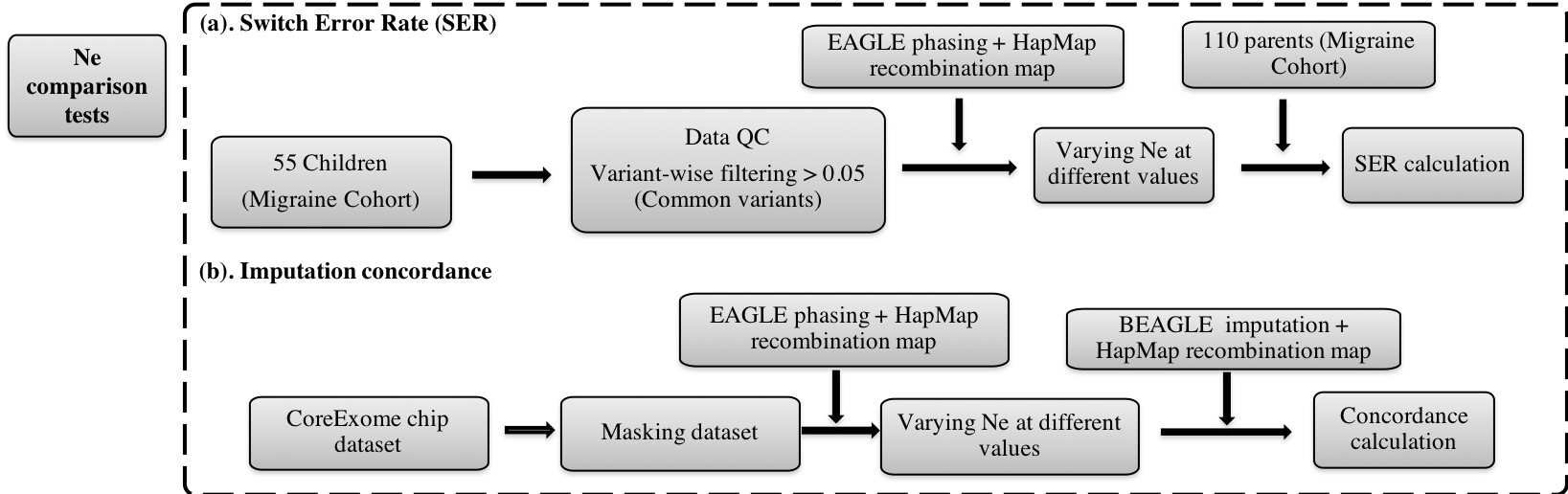

**Figure S3**: Flowchart for effective population size (Ne) comparison tests

To test, whether varying Ne has an effect among the studied populations in downstream analyses (phasing and imputation accuracy). We found that the average switch error rates (SER, see Supplementary Methods for description) was similar across all the tested values of Ne across chromosomes (1.9-2.7%, Figure S4). The average imputation concordance for variants in the >5% MAF (minor allele frequency) group was ~98%, with a drop in concordance rates towards the low frequency (1-5% MAF, 90-95%, Figure S5) and rare (< 5% MAF, 76-87%, Figure S5) frequency variant groups.

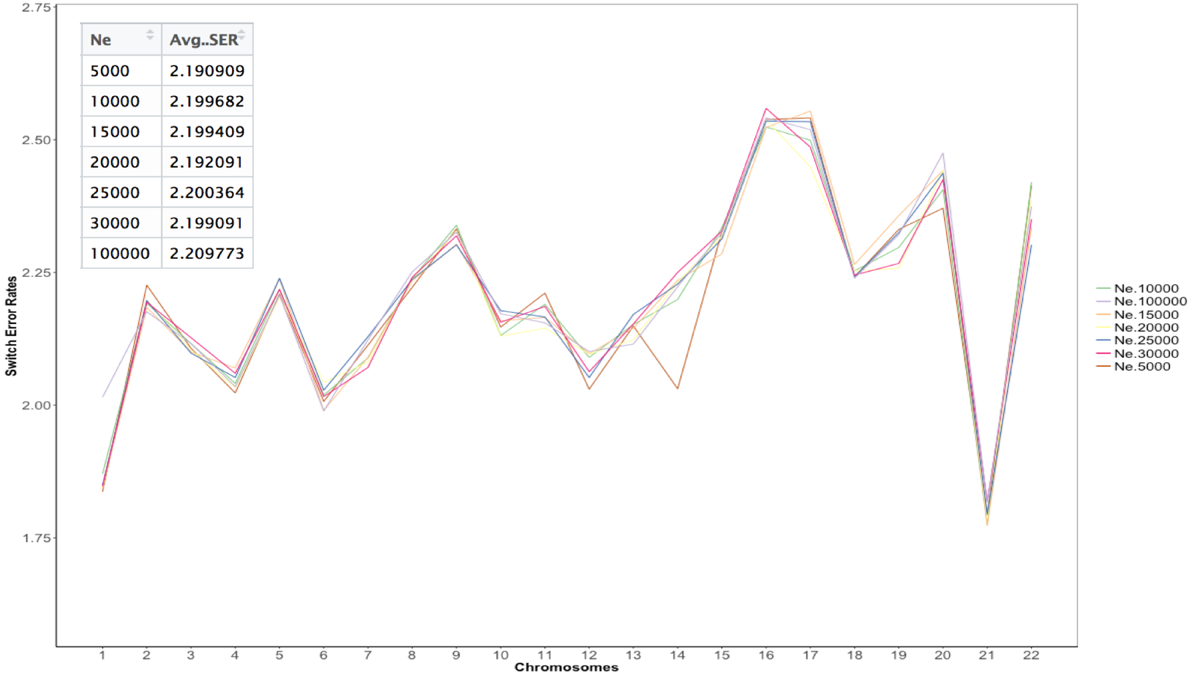

**Figure S4**: Comparison of Switch Error Rates across all autosomes for different values of Effective Population Size (Ne).

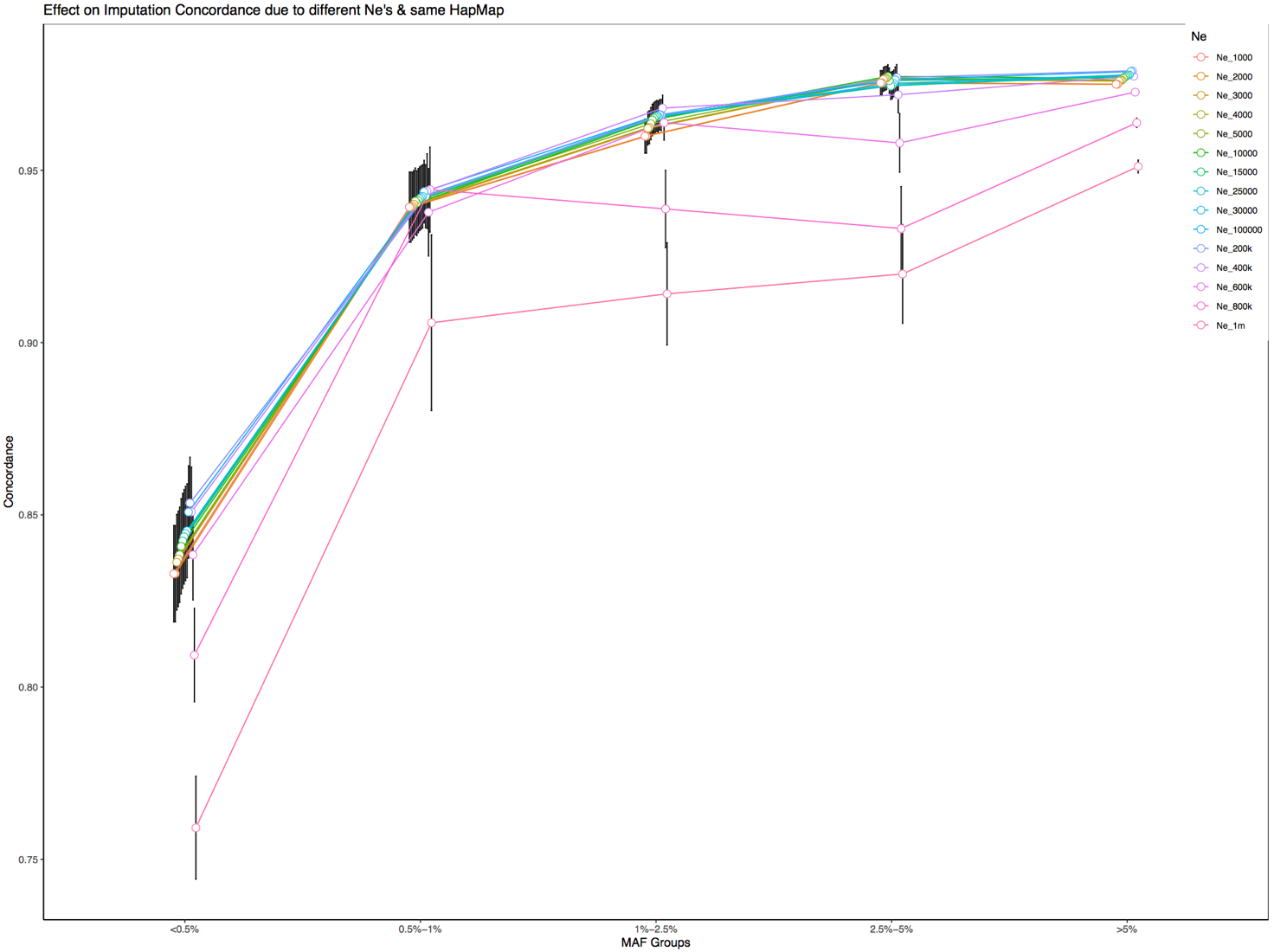

**Figure S5:** Comparison of Imputation Concordance across different Minor Allele Frequency (MAF) groups for a range of Effective Population Sizes (Ne) (Ne = 1000, 2000, 3000, 4000, 5000, 10000, 15000, 25000, 30000, 100000, 200000, 400000, 500000, 800000, 1000000)

**Effect of Reference Panel size on the downstream analyses**

To check whether reference panels used during haplotype phasing and imputation made any impact on the switch error rates and imputation concordance respectively, we did an additional step of comparison by randomly choosing 500 individuals from the previously used (n=2690) Finnish reference dataset.

The smaller reference panel had higher switch error rates (SER) when compared to the larger sized reference panel (Figure S6) but had lower values when compared to the setting which didn’t use any reference panel for phasing.

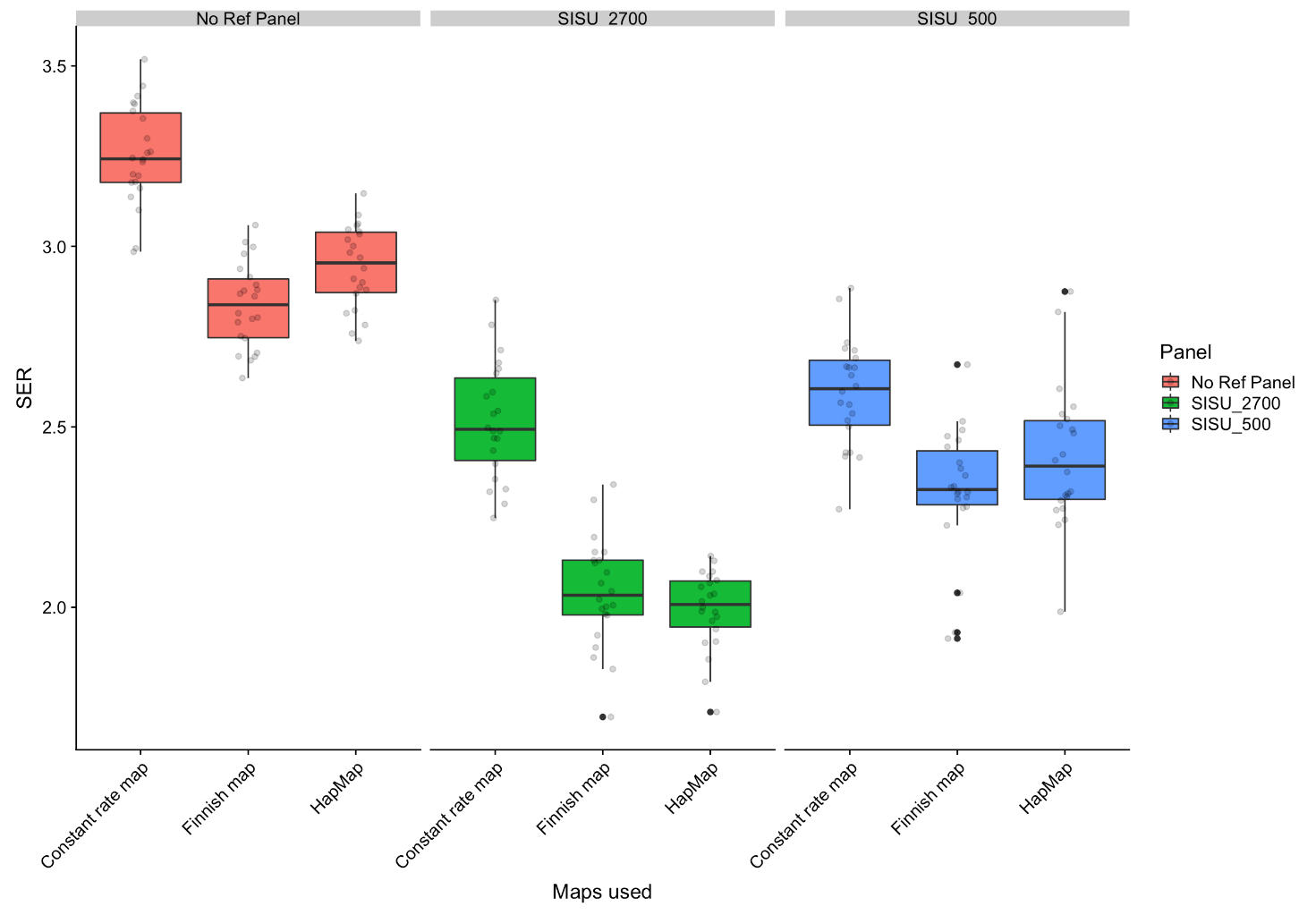

**Figure S6:** Comparison of Switch Error rates across the different reference panel groups under different recombination map conditions

We found that the smaller panel (n=500) yielded lower imputation accuracy (concordance rate 72~76%, Figure S7). The concordance rate was lower when the test dataset was

phased without reference panels, with larger Finnish panel (n= 2700) yielding higher values (concordance rate 72~77%, Figure S8) than the smaller panel (concordance rate 52~57%, Figure S8). This highlights the role of sample size of the reference panels used.

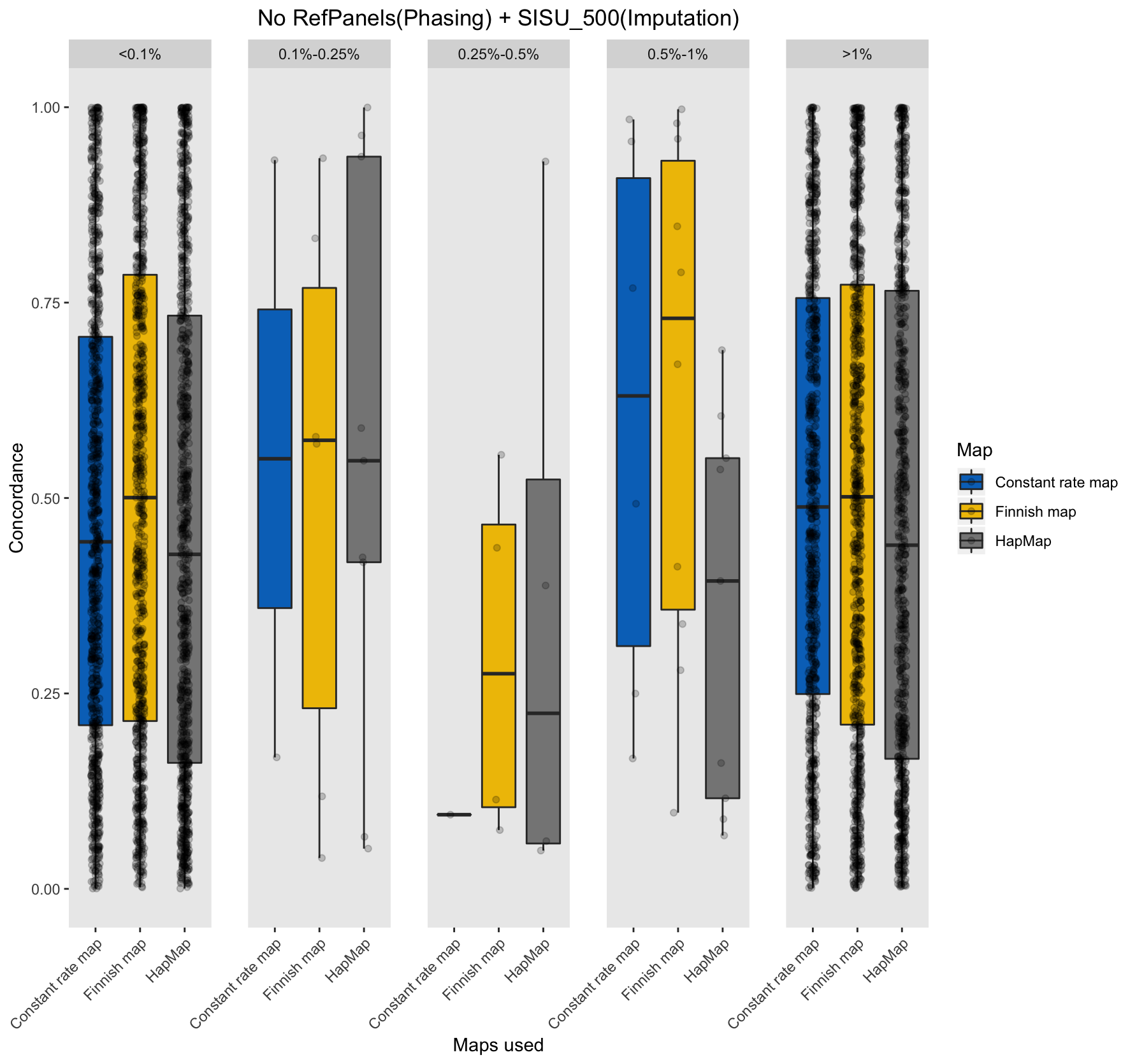

**Figure S7:** Comparison of Imputation Concordance across different Minor Allele Frequency (MAF) groups for a range of different recombination map combinations phased with NO reference panels

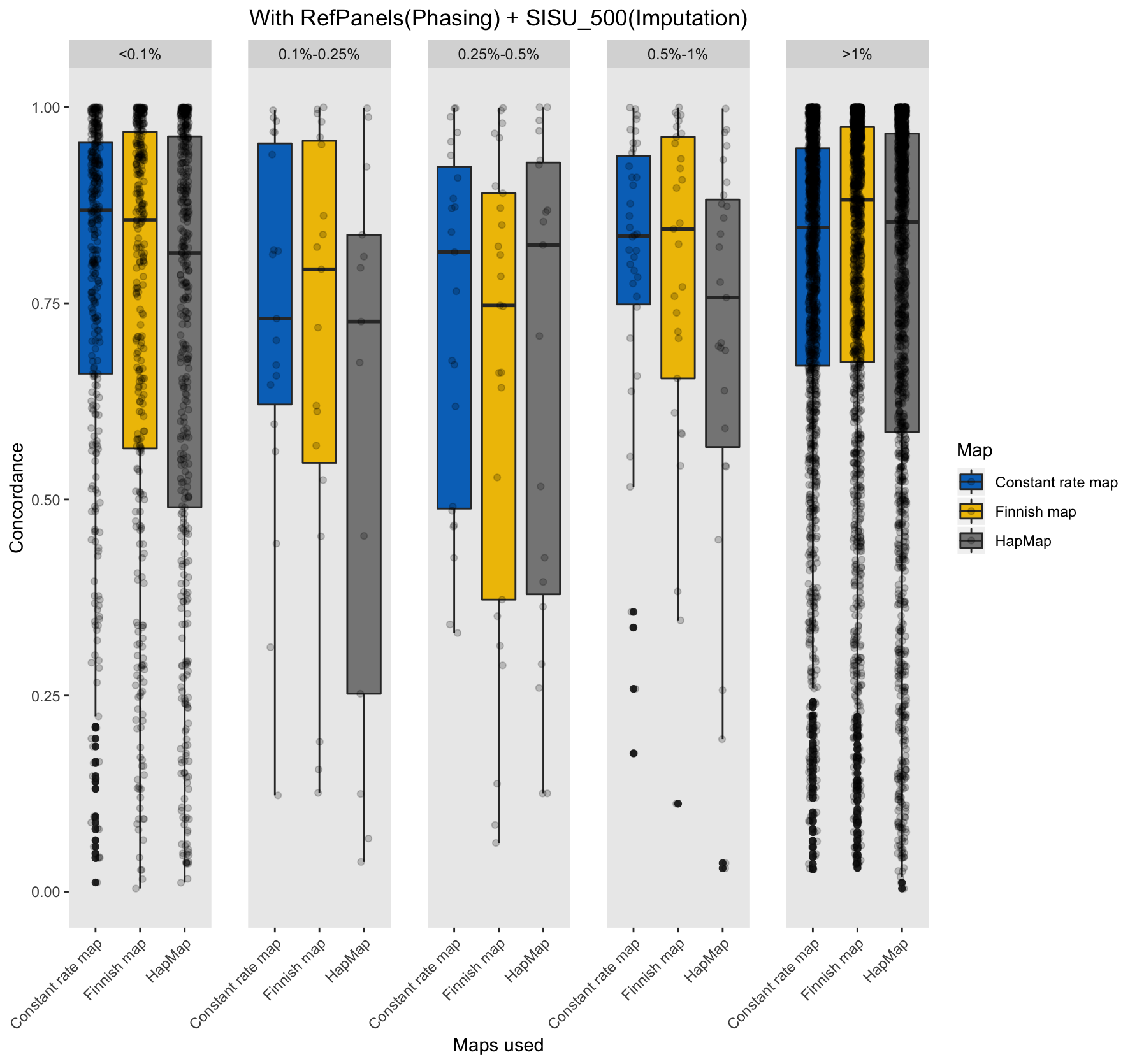

**Figure S8**: Comparison of Imputation Concordance across different Minor Allele Frequency (MAF) groups for a range of different recombination map combinations phased with reference panels
